# Real time mobilization of a novel diatom *Mutator-Like Element* (MULE) transposon to inactivate the uridine monophosphate synthase (UMPS) locus in *Phaeodactylum tricornutum*

**DOI:** 10.1101/2023.01.02.522487

**Authors:** Raffaela M. Abbriano, Jestin George, Tim Kahlke, Audrey S. Commault, Michele Fabris

## Abstract

Diatoms are photosynthetic unicellular microalgae that drive global ecological phenomena in the biosphere and are emerging sustainable feedstock for an increasing number of industrial applications. Diatoms exhibit enormous taxonomic and genetic diversity, which often result in peculiar biochemical and biological traits. Transposable elements (TE) represent a substantial portion of diatom genomes and have been hypothesized to exert a relevant role in enriching genetic diversity and centrally contribute to genome evolution. Here, through long-read whole genome sequencing, we identified a novel Mutator-Like Element (MULE) in the model diatom *Phaeodactylum tricornutum,* and we report the direct observation of its mobilization within the course of one single laboratory experiment. Under selective conditions, this novel TE inactivated the *uridine monophosphate synthase* (*UMPS*) gene *of P. tricornutum,* one of the two only endogenous genetic loci currently targeted for selectable auxotrophy in functional genetics and genome editing applications.

We report the first, real-time observation of the mobilization of a transposon in diatoms that possesses novel peculiar features. These include the combined presence of a MULE transposase domain with Zinc finger, SWIM-type domains, and of a diatom-specific E3 ubiquitin ligase of the zinc finger UBR type, which indicate a novel mobilization mechanism. Our findings provide new elements for the understanding of the role of TEs in diatom genome evolution and in the enrichment of intraspecific genetic variability. Ultimately, this raises relevant concerns on the targeting of loci such as *UMPS* as selectable markers for functional genetics and biotechnological applications in diatoms.

**Significance Statement:** We identified a novel DNA transposon in the diatom *Phaeodactylum tricornutum*. This new Mutator-Like Element encodes a transposase and a diatom-specific E3 ubiquitin ligase, which suggest a novel mobilization mechanism. We documented independent insertions in real-time, which spontaneously inactivated the *uridine monophosphate synthase* (*UMPS*) locus, a common selectable marker. We provide new insights on the role of transposons in diatom genome dynamics and evolution and on the unsuitability of *UMPS* as selection locus in diatoms.

## Introduction

Diatoms are unicellular photosynthetic microalgae with central ecological roles and at the centre of increasing biotechnological interest. They are widespread among the planet’s diverse aquatic and oceanic environments due to their unique and efficient metabolic traits. As such, diatoms are among the most relevant primary producers and main drivers of oceanic geochemical cycles, in addition to being a major component of aquatic food-chains (Armbrust, 2009). A number of diatom species also find industrial and commercial applications related to their biochemical traits, including aquaculture and the photosynthetic production of high-value bio-products, such as polyunsaturated fatty acids, pigments, and proteins (Butler *et al*., 2020; Fabris, Abbriano, *et al*., 2020). In particular, the genetically tractable *Phaeodactylum tricornutum* is a model organism for advanced bioengineering applications, including metabolic engineering and synthetic biology approaches, that could expand the biosynthetic capacity of diatoms beyond their natural traits (Hempel, Bozarth, *et al*., 2011; Hempel, Lau, *et al*., 2011; D’Adamo *et al*., 2018; Fabris, George, *et al*., 2020; Pampuch *et al*., 2022; Slattery *et al*., 2022; Windhagauer *et al*., 2022).

The unique metabolic capacities of diatoms have originated through a peculiar evolutionary path, which included at least two independent endosymbiotic events, and numerous horizontal gene transfers (HGTs) (Vancaester *et al*., 2020). Through these, diatoms have acquired genes from all domains of life (Bowler *et al*., 2008). These genetic features have then been reshuffled and reshaped in a wealth of configurations, and resulted in their enormous genetic and taxonomic diversity, by mechanisms that are still largely unknown.

A major driver of genetic diversity in diatoms, as in other organisms, is the presence and action of transposable elements (TE) (Maumus *et al*., 2009). In several eukaryotes, TEs have been shown to be widespread and to undergo frequent mobilization and rearrangement, particularly during stress, and have been hypothesized to play central roles in genome evolution (Sun *et al*., 2020). Recent analyses highlighted that the portion of *P. tricornutum* genome composed by TEs is much larger than previously thought and suggested that the effect and influence of TEs in diatoms is a key evolutionary factor in the wild (Filloramo *et al*., 2021).

Even in controlled laboratory conditions, diatoms can accumulate high haplotype diversity even within the same clonal cultures, where genotype diversity is enhanced by frequent interhomologous recombination events, which are favoured in stress conditions (Bulankova *et al*., 2021). Aspects of genome stability in diatoms are important to understand their evolution and role in the ecosystems, but also in assessing the robustness of genetically engineered strains. In this latter scenario, many applications are based on the artificial perturbation of specific small portions of their genomes, such as in the case of the maintenance of functional selectable marker or transgenes, or both.

The enzyme uridine monophosphate synthase (UMPS) is involved in *de novo* pyrimidine biosynthesis and its genetic locus is a relevant selectable marker in diatoms (Sakaguchi *et al*., 2011; Serif *et al*., 2018; Pampuch *et al*., 2022). UMPS knock-out mutants are tolerant to 5-fluoroorotic acid (5-FOA) and are uracil auxotrophic. 5-FOA is a chemical analogue of oroate, which is converted by UMPS to form orotidine-5’-phosphate (EC 2.4.2.10), which is then converted by UMPS again to form uridine-5’-monophosphate (EC 4.1.1.23), a precursor of uracil (Fig. 1b). In the presence of 5-FOA and uracil, UMPS catabolises the 5-FOA analogue compound to generate 5-fluorouracil (5-FU), a toxic molecule that causes cell death (Fig.1a). When UMPS is inactivated, the resultant mutant requires uracil supplementation in the medium, but is also unable to catabolise 5-FOA into toxic 5-FU, making this a useful selectable marker gene for screening microbes on 5-FOA selection plates (Fig. 1a). This marker, along with an alternative strategy that targets the adenine phosphoribosyl transferase (APT), currently represents the only approaches available for ribonucleoprotein (RNP) based CRISPR applications in diatoms (Serif *et al*., 2018; Slattery *et al*., 2018). For this locus to be a reliable genetic marker, it needs to be particularly stable. However, numerous genome editing experiments carried out in our laboratory targeting *PtUMPS* suggested that this locus in *P. tricornutum* (*Phatr3_J11740*) is particularly prone to produce false positives, as we consistently observed the emergence of diatom colonies able to grow on 5-FOA without their *PtUMPS* locus being targeted by nuclease. To date, there are no reports of 5-FOA induced mutation in diatoms. However, it has previously been reported that wild type *P. tricornutum* could become reliant on uracil and resistant to 5-FOA (RURF) following chemical mutagenesis using N-ethyl-N-nitrosourea (ENU) and selection using 100 - 300 μg/mL of 5-FOA (Sakaguchi *et al*., 2011).

**Figure 1.**
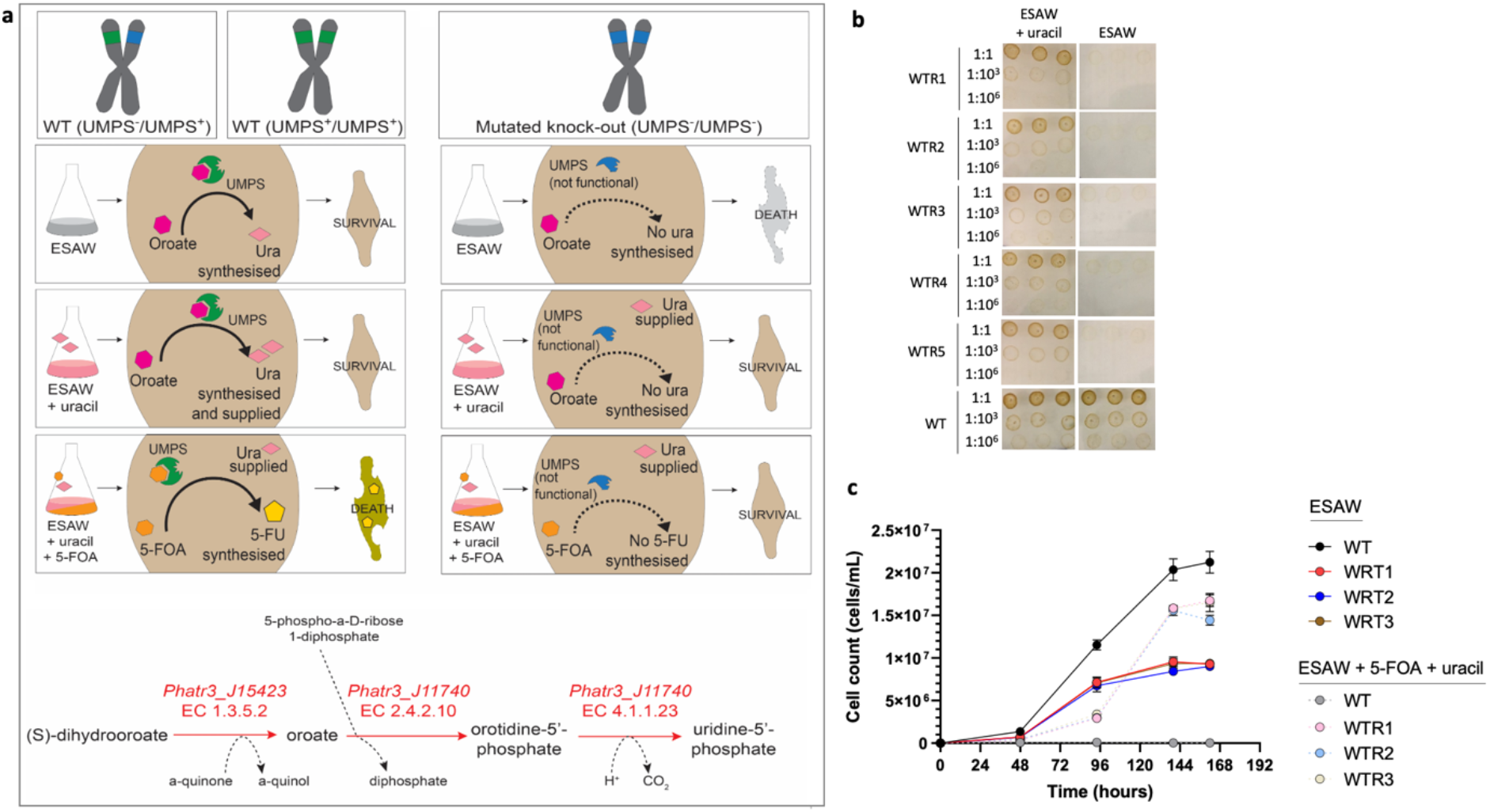
Phenotyping of 5-FOA resistant WT *P. tricornutum* strains (WTR). a) Schematic representation of WT and UMPS-deficient phenotypes either in presence or in absence of 5-FOA and uracil; b) uracil auxotrophy assessed by spotting either WTR or WT diatoms on ESAW-agar medium supplemented with uracil 50 μg/mL or ESAW-agar medium; c) growth of three WTR strains compared to WT controls cultivated in liquid ESAW medium in presence or absence of 5-FOA (300 μg/mL) and uracil 50 μg/mL (n=3, error bars represent standard error).

Here, we isolated and characterized *P. tricornutum* cell lines that spontaneously emerged as RURF-like in the absence of exogenous genetic manipulations and known mutagens. We provide phenotypic and genotypic evidence of the systematic and reproducible disruption of the *PtUMPS* locus through the activation of a novel transposable element mobilized during 5-FOA selection.

## RESULTS AND DISCUSSION

### Wild type *P. tricornutum* exposed to 5-FOA produces colonies with stable RURF-like phenotype

To investigate the mechanism of occurrence of the 5-FOA-resistant diatom phenotype, we subjected wild type (WT) *P. tricornutum* cultures (1.5·10^8^ cells) to a range of concentrations of 5-FOA (0-1000 μg/mL), including those (100-300 μg/mL) typically used in CRISPR-Cas9 RNP-mediated experiments (Serif *et al*., 2018), and supplemented it with 50 μg/mL uracil to rescue the possible inactivation of *PtUMPS*. After seven weeks, hundreds of 5-FOA resistant colonies (WTRs) emerged on plates with 5-FOA concentrations ranging from 50-150 μg/mL, several colonies were found on plates with 5-FOA 300 μg/mL (Fig. S1), while no diatoms survived higher concentrations (600-1000 μg/mL, Fig. S1). We hypothesized that the WTR strains generated in this study had the same RURF phenotype described by Sakaguchi et al. (2011).

To characterize the WTR phenotype, we selected five random colonies that originated from five different plates. To confirm the auxotrophy of WTR cell lines for uracil, we grew them in ESAW without uracil for one week to deplete their endogenous cellular uracil pools and then dilution plated them onto ESAW or ESAW supplemented with uracil. Whereas the WT lines were able to grow on ESAW plates without uracil supplementation, none of the WTR cell lines were able to survive. However, the WTR phenotype could be rescued by the presence of uracil in the medium (Fig. 1b). To further confirm the phenotype, three WTR primary colonies and one untreated WT colonies were grown in the absence and presence of 300 μg/mL 5-FOA and 50 μg/mL uracil over seven days (Fig. 1c). As expected, untreated WT diatoms grew normally in ESAW medium, indicating the presence of a functional PtUMPS enzyme and endogenous uracil biosynthesis, whereas when cultured in presence of 5-FOA and uracil, no growth was observed, due to the functional PtUMPS enzyme metabolizing 5-FOA into toxic 5-FU. In absence of 5-FOA and uracil, WTR diatoms grew at a reduced rate in the early exponential phase, from 48 hrs to 96 hours, most likely due to the declining presence of an intracellular uracil pool available to the cells for RNA biosynthesis. Cell growth drastically slowed after 96 hours and eventually arrested at approximately 144 hours, suggesting that the uracil reserves were depleted. In contrast, these strains were all able to grow in the presence of 5-FOA and uracil, confirming that they were unable to metabolize 5-FOA into toxic 5-FU, but able to use supplemented uracil for RNA biosynthesis (Fig. 1c).

Together, these results confirmed that 5-FOA exposure results in a RURF-like phenotype (Sakaguchi et al., 2011) in WTR diatoms and strongly suggested that the PtUMPS enzyme might have been inactivated in these cell lines.

### WTR diatoms cell lines harbour large mutations in the *PtUMPS* locus

To determine whether the inactivation of PtUMPS in WTR lines was due to genetic disruption of the *PtUMPS* gene, we interrogated this locus using PCR and Sanger sequencing in the selected five WTR and in one WT cell lines (Fig 1b). While we were able to amplify the full *PtUMPS* locus from genomic DNA of untreated WT diatoms, we were not able to amplify it from any of the WTR strains except from WTR4 (Fig. 2a). Next, we used different primer combinations that spanned all three exons of *PtUMPS* (Fig. 2b, Table S1) to try to detect disruptions at the origin of the *PtUMPS* inactivation. We obtained correctly amplified PCR products from untreated WT samples using all four primer combinations (Fig. 2a). However, in WTR samples we could only amplify portions of *PtUMPS* when primers were designed to anneal in the first two exons. We were unable to amplify the terminal region in all WTR samples, with the only exception of WTR4, suggesting the disruption of the *PtUMPS* gene in the remaining WTR cell lines. The PCR results indicate the presence of a large insertion or deletion (INDELs) in all WTR cell lines, except for WTR4 (Fig 2c), which nonetheless clearly showed a phenotype associated with a non-functional *PtUMPS*. Sanger sequencing of the full *PtUMPS* gene from the untreated WT sample revealed a heterozygous genotype consisting of the same 16 single nucleotide polymorphisms (SNPs) identified by Sakaguchi et al. (SNP-1–16), as well as an additional three SNPs (SNP-A–C) (Fig. S2). These were identified by the presence of peak doublets in the sequencing chromatogram (Fig. S2) and suggested that untreated WT *P. tricornutum* harboured one functional *PtUMPS* allele (allele A) and a second non-functional allele (allele B, Fig 1a). Therefore, we hypothesized that WTR mutants contained only the non-functional allele B. Given that we were only able to amplify the 5’ end of *UMPS* in the WTR mutants, we Sanger sequenced amplicon 40-74 from WTR1, WTR2, and WTR3 (Fig. 2c). Interestingly, the sequences obtained for amplicon 40-74 matched functional allele A (Fig. S2). The obtained WTR genotype differed from the predicted *PtUMPS* WT sequence and putatively encoded a 254 amino acids long protein (instead of the 518 amino acids of the full-length protein) without the purine/pyrimidine phosphoribosyl transferase (PPRT) active domain, and with a single point mutation in the coding sequence causing the substitution of a methionine by an isoleucine (Fig. 2c).

**Figure 2.**
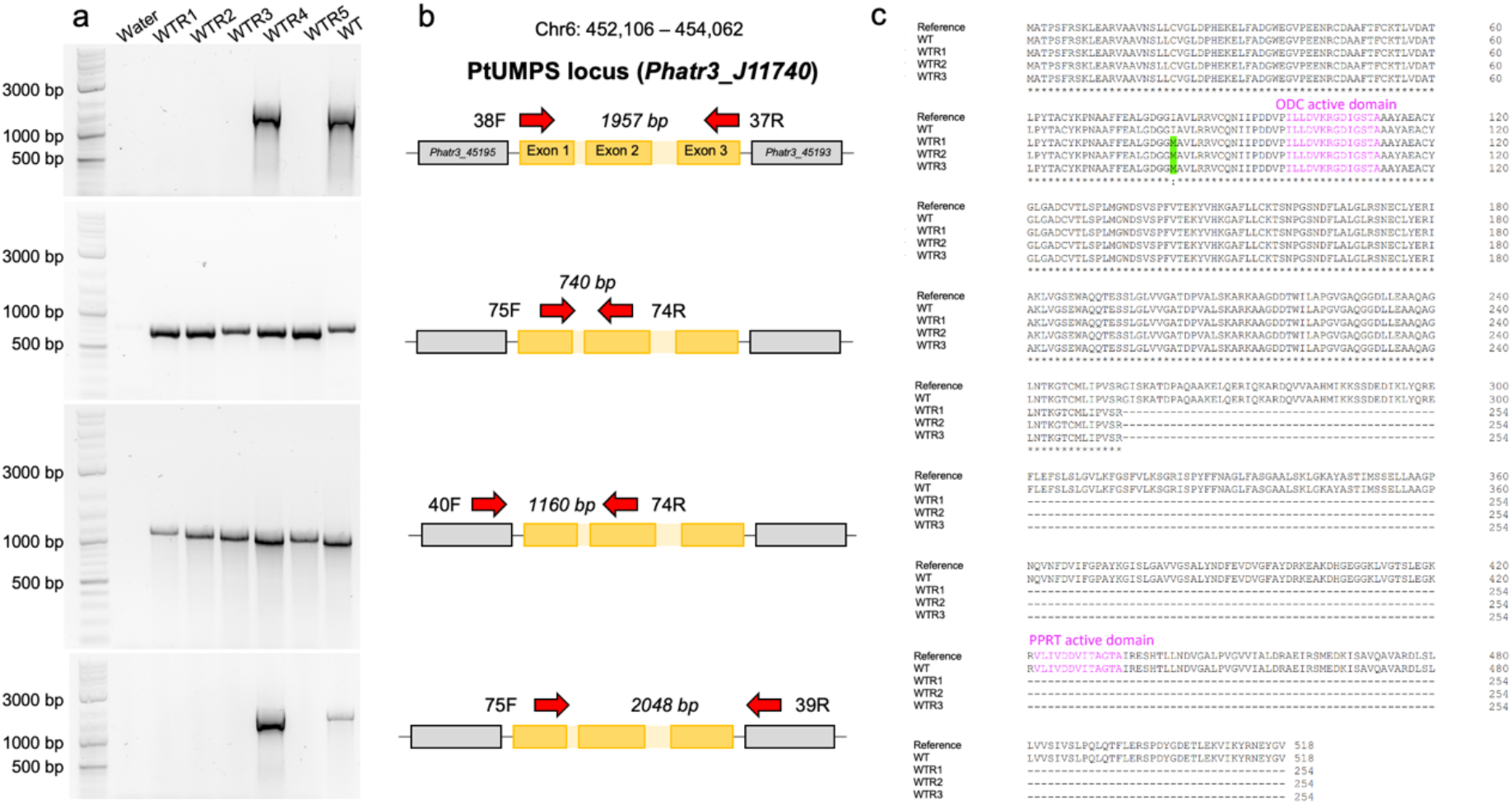
Genotyping of 5-FOA resistant WT *P. tricornutum* strains (WTR). a) PCR amplification of the *PtUMPS* locus in WTR and WT strains, with different primers combinations. b) Schematic representation of the amplified genomic regions and expected amplicon size in case of intact *PtUMPS* locus (in reversed configuration for ease of interpretation). c) Multiple sequence alignment of PtUMPS (Uniprot Accession C6L824) protein to translated nucleotide sequences obtained from the *PtUMPS* amplicons of WT, WTR1, WTR2, WTR3 obtained in this study. All three WTR strains demonstrate a single point mutation (green) causing an amino acid change from isoleucine to methionine and stunted coding region as well as a truncated protein (254 amino acids instead of the full 518 amino acids), and the loss of the PPRT active domain (pink).

Considering these results, we rejected our original hypothesis that the RURF phenotype in *P. tricornutum* WTR strains was caused by point mutations in the functional allele, as reported by Sakaguchi et al. (2011), but we hypothesized instead that the phenotype could originate from a larger chromosomal mutation which caused the severe truncation of the *PtUMPS* gene and the loss of the PPRT active domain. Such large chromosomal mutations have indeed been reported in other species, such as the loss of entire chromosomes following 5-FOA exposure in *Candida albicans* (Wang *et al*., 2004; Wellington and Rustchenko, 2005; Wellington *et al*., 2006).

### A novel *mutator-like element* (MULE) transposon inactivated the *PtUMPS* locus in WTR diatoms

To test for the presence of large re-arrangements affecting the *PtUMPS* locus in WTR diatom strains, we employed Oxford Nanopore long-read sequencing and assembled the genomes of all WTR cell lines as well as of the WT control (see Table S2 and S3 for genome sequencing and assembly statistics, respectively). Investigation of the *PtUMPS* locus using the Integrated Genomics Viewer (Robinson *et al*., 2011) revealed mutations of the *PtUMPS* gene in all WTR cell lines, while the corresponding WT locus was found to be intact. Interestingly, all cell lines except for WTR4 presented a single large insertion ranging from 2.3 to 3.7 kb in the *PtUMPS* coding sequences. These were confirmed by analysing the assembly with the structural variant caller Sniffles (Sedlazech *et al*., 2018) using the raw sequencing reads mapped against the published *P. tricornutum* genome. In contrast to the other WTR samples, WTR4 only presented a small deletion (15 bp) in exon 1 (Fig 3). All mutations in the *PtUMPS* locus of WTR cell lines resulted in a disrupted gene sequence, corroborating the hypothesis that the observed phenotypes could be attributed to *PtUMPS* inactivation by larger-scale chromosomal modifications, as suggested by the PCR-based genotyping (Fig 2).

**Fig. 3.**
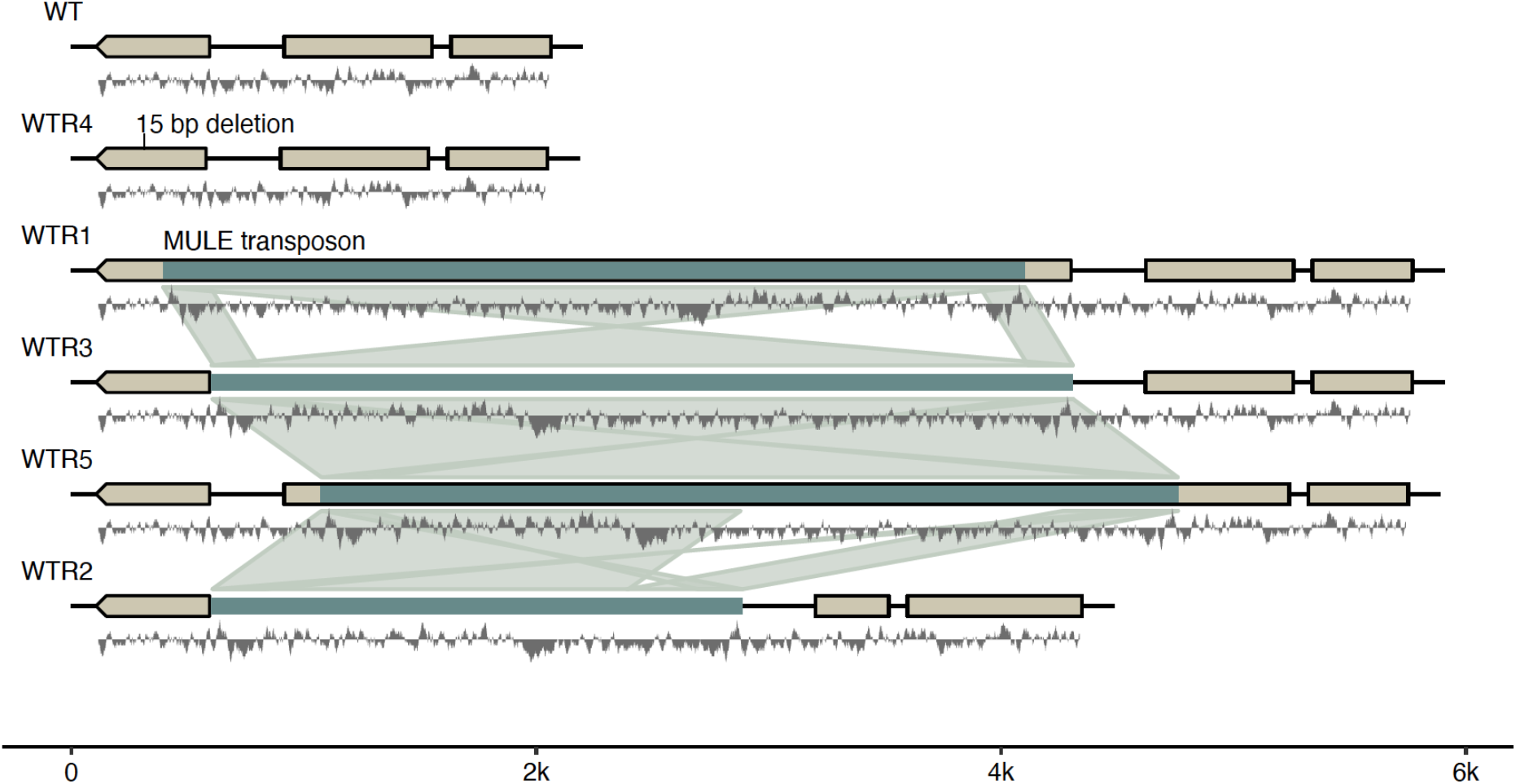
Schematic representation of the genetic re-arrangements in the *PtUMPS* locus. Assembled genome sequences at the *PtUMPS* locus, in WT and WTR lines. The coding sequence of the *PtUMPS* gene is shown in beige (reverse orientation). Teal blocks represent insertion sequences annotated MULE transposons, and light teal ribbons represent synteny links between insertions at a cutoff of >80% nucleotide sequence similarity. A GC track calculated with a 20 bp sliding window with a midline of 50% GC content is shown in grey beneath each sequence.

Further investigation of the insertion sequences at the *PtUMPS* locus of cell lines WTR1, 2, 3, and 5 revealed the presence of *terminal inverted repeats* (TIR) of approximately 200 bp at the beginning and end of the insertions as well as a high degree of overall sequence similarity (Fig 4a, S3) indicating the insertion of a transposable element into the *PtUMPS* region of these cell lines. Sequence analysis around the insertion sites also revealed short (9 bp) target site duplications (TSDs) that externally flanked the TIRs (Fig 4a, Table S4). TSDs delimit the transposable element sequence as molecular ‘scars’ of the TE insertion (Fig. 4a, Fig S3), and are distinctive features of MULEs (Wicker *et al*., 1989). The similar nature and low GC content of the TSDs (Table S4) suggests that TE insertion may be specific to TA-rich regions.

**Fig. 4.**
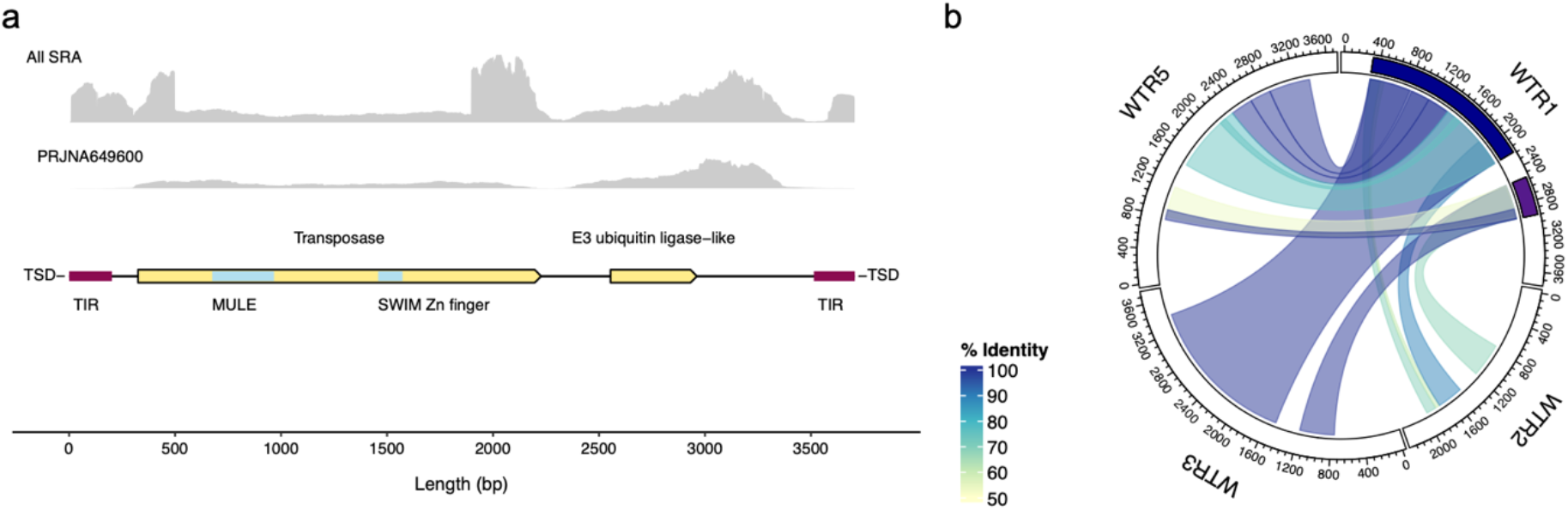
Structure and features of the TE which disrupted *PtUMPS* in WTR1. a) Full length (3707 bp) of the entire transposon sequence in WTR1. ORFs corresponding to the transposase and the E3 ubiquitin ligase-like protein are depicted in yellow, with predicted MULE and SWIM Zn finger domains in blue. The TIR sequences are represented in maroon and flanked by 9 bp TSD sequences. The RNAseq data is from available reads in NCBI SRA (top) and from samples PRJNA649600; b) Conservation of structure and features of the TE from WTR1 (dark blue) in WTR2, WTR3 and WTR4, WTR1.

Gene prediction and functional annotation of the TE sequences using InterProScan (Jones *et al*., 2014) revealed the presence of an open reading frame (ORF) putatively encoding a protein with a MULE transposase domain and a Zinc finger, SWIM-type domain (Fig. 4) in WTR1, and WTR3. Analysis of the TE sequences of WTR2, and WTR5 showed multiple mutations and frame-shifts possibly disrupting the predicted transposase ORF in WTR5 as well as a large deletion in the WTR2 TE sequence, resulting in the deletion of most of the transposase ORF (Fig 4 b, Table S5). We identified multiple orthologs of the putative MULE transposase harboured in the TE sequence, in *P. tricornutum (Phatr3_EG01202, Phatr3_EG00036, Phatr3_EG00117, Phatr3_EG02053, Phatr3_J33421, Phatr3_EG01875, Phatr3_EG00351, Phatr3_J48581),* which show similarities to other diatom predicted proteins that contain MULE transposase domains and, to a lesser extent, to hypothetical proteins in eudicots that do not contain it. Interestingly, InterProScan analyses of the inserted TE sequences revealed the presence of the *Phatr3_J50367* gene, either in full or partial configuration. Searches on NCBI BLASTp and Plaza Diatoms (Vandepoele *et al*., 2013), revealed that this gene possess three homologues with high similarity in *P. tricornutum (Phatr3_EG02053, Phatr3_J48581* and *Phatr3_EG02300*). *Phatr3_J50367* belongs to gene family specific to raphid diatoms, with no significant similarities outside this group, and putatively encodes an E3 ubiquitin ligase of the zinc finger UBR type (Fig. 4a), according to EggNOG predictions (Huerta-Cepas *et al*., 2019) reported in Plaza Diatoms. Similar to the MULE transposase, the complete sequence of this ORF was only found in some of the inserts. We detected the full sequence of *Phatr3_J50367* in WTR1 and WTR3 but only partial sequences in the other samples (Fig. 4b, Table S2).

To assess whether both the ORFs encoding the transposases and the E3 ubiquitin ligases are actively transcribed in *P. tricornutum* and not pseudogenes, we searched for evidence of their expression in publicly available transcriptomics data. Manual inspection of the mapped reads using the Integrated Genome Viewer (Robinson et al., 2011) showed evidence of the expression of the complete transposase as well as large parts of the transposon (Fig. 4a). Interestingly, expression of the transposon was particularly pronounced in transgenic *P. tricornutum* cell lines overexpressing a *Thalassiosira pseudonana* chitin synthase (SRA project #PRJNA649600).

The presence of a MULE transposase in the TE sequence ruled out the possibility that the TE we identified falls into the sub-class of Pack-MULE TEs, which instead are “not autonomous” as they do not include a transposase and generally carry portions of several other host genes within them (Dupeyron *et al*., 2019). The presence of UBR type zinc finger E3 ubiquitin ligases in some of the TE sequences suggests that mobilization of this MULE transposon mechanism might involve an active post-translational regulation, based on protein degradation, which has never been described before. TE regulation mechanisms involving protein ubiquitination and histone binding in *Arabidopsis thaliana* have been reported (Kabelitz *et al*., 2016). While negative regulatory roles of E3 ubiquitin ligases have been observed in LTR retrotransposons in mice (Maclennan *et al*., 2017), they have never been reported in the case of Class 2 DNA transposon of the MULE type. Given the lack of homologous of *Phatr3_J50367* outside the diatom taxonomic group, we hypothesize that this novel mechanism of DNA transposition might be diatom-specific. Inserts detected in WTR1, 3 and 5 exhibited the highest similarity, while the one found in WTR2, although still highly similar, lacks a large portion of the sequence present in the other samples (Fi. 4b, Table S5). This suggests that there may be some variation in the insert length or modifications to the inserted sequence might occur during the insertion process.

In contrast to WTR1, WTR4 and WTR5, the presence of a small 15 bp deletion in the *PtUMPS* locus of WTR4 is likely to have emerged independently and subsequently selected by the experimental conditions, in agreement with the uracil auxotrophy phenotype and the PCR amplification of the exons (Fig. 2). This suggests that selection using 300 μg/mL 5-FOA can induce presumably chemically-mediated mutations in *P. tricornutum* CCAP 1055/1, resulting in RURF phenotypes.

### The activation of the novel MULE transposon is favoured by 5-FOA selection

The fact that we observed interruption of the *PtUMPS* locus in all five WTR mutant cell lines suggest that selection with 300 μg/ml 5-FOA clearly creates a strong bias towards the occurrence and selection of *PtUMPS* mutations. In four out of the five mutants we analysed, the *PtUMPS* locus was disrupted by the same MULE transposon that was not present in the same location in the parental cell line before the experiment (Fig. 3), suggesting that our selection conditions favoured TE-mediated mutations over TE-independent ones (WTR4) or that TE-mediated mutations are more frequent and/or more effective. In addition, the high degree of similarity between the TIR sequences, often used to estimate the age of the element (Hanada *et al*., 2009), indicates that transposons mobilized recently and that they originated from the same genomic source. Together, this provides clear evidence that in our experiments *PtUMPS* was disrupted by a TE, which was activated during the selection process, potentially triggered by sudden stress conditions represented by the accumulation of the toxic 5-FU. Transposons and MULEs in particular are known to mobilize as a response to certain stress conditions in several organisms, either as an active coordinated response, or as epigenetic de-regulation (Negi et al., 2016).

We further explored whether the MULE-like TE was equally active in the genome of cell lines where it directly inactivated *PtUMPS* (WTR1-3 and WTR5), in WTR4 where *PtUMPS* was disrupted by a TE-independent mutation, and in WT where *PtUMPS* was left intact. Using the transposon that disrupted the *PtUMPS* locus in WTR1 as query, we searched for highly similar regions with strict parameters (length >1.5 Kb, lowest similarity 80% and E-value 0.0) throughout the genomic assemblies we generated (with genome coverages ranging 18x to 68x), as well as in the publicly available one (Rastogi et al., 2018). In untreated WT, this specific MULE transposon was found to occur in only 5 instances, which is comparable to the frequencies detected in publicly available genome assemblies. In contrast, the transposon was detected in much higher frequencies ranging from 20 to 131 copies (Table S2) in WTR cell lines subjected to 5-FOA selection. Interestingly, the WTR4 cell line that developed a TE-independent *PtUMPS* mutation exhibited a MULE TE mobilization frequency (25 copies) that is comparable to that of TE-dependent cell lines.

Although this does not provide evidence of the targeted mobilization of the MULE TE to the *PtUMPS* locus in direct response to 5-FOA selection, these results demonstrate that the observed MULE-mediated mutation occurred within the timeframe of the experiment and suggest that MULEs may be activated during cellular stress, either as regulated mechanism or as consequence of de-repression.

### Implications of the use of 5-FOA as selection agent in diatoms

The mutant cell line WTR4 resulted from a TE-independent 15 bp deletion in the *PtUMPS* locus, suggesting that selection using 300 μg/mL 5-FOA can induce presumably chemically-mediated mutations in *P. tricornutum* CCAP 1055/1 that result in RURF phenotypes. This phenotype has previously been obtained in *P. tricornutum* strain UTEX LB 642 following chemical mutagenesis using N-ethyl-N-nitrosourea and selection with 5-FOA (Sakaguchi *et al*., 2011), and in *P. tricornutum* strain CCAP1055/1 following targeted gene knock-out of *UMPS* gene via CRISPR-Cas9 gene editing (Serif et al., 2018; Slattery et al., 2020). In these works, diatoms were selected on 100 μg/mL 5-FOA. Our findings demonstrate that it is possible that 5-FOA can be mutagenic in *P. tricornutum* at concentrations above 100 μg/mL. These results also suggest that 5-FOA may be inappropriate for selection of transformants or genome edited diatoms, as chemical and TE-induced mutagenesis cannot be excluded. 5-FOA has shown mixed mutagenic effects in different organisms, including *S. cerevisiae* (Klein *et al*., 2014; Backhaus *et al*., 2017; Scott *et al*., 2018) and others, such as *Y. lipolytica* (Lv *et al*., 2019), as well as *E. coli* (Standage-Beier *et al*., 2015; Brandsen *et al*., 2018). However, 5-FOA is highly mutagenic in *C. albicans* (Wang *et al*., 2004; Wellington and Rustchenko, 2005; Wellington and Rustchenko, 2005). There is also a report of 5-FOA inducing mutation in *Sulfolobus acidocaldarius* (Kondo *et al*., 1991) and in *S. cerevisiae* (Hao *et al*., 2016). Furthermore, 5-FOA has been used for generating spontaneous RURF phenotypes in the dinoflagellate *Symbiodinium* SSB01 following exposure to 200 μg/mL 5-FOA (Ishii *et al*., 2018), and the red alga *Cyanidioschyzon merolae* 10D following treatment with 800 μg/mL 5-FOA (Minoda *et al*., 2004).

### Implications of the mobilization of TE-mediated, non-homologous genome rearrangements in diatom genome evolution and biotechnology

Recently, it has been reported that diatoms use a yet elusive mechanism of mitotic interhomolog recombination to increase phenotypic plasticity, which is particularly enhanced when diatoms face sudden environmental stress, and this phenomenon has been observed to occur specifically at mutated *PtUMPS* loci to restore the WT allele (Bulankova *et al*., 2021). Here we provided evidence of the mobilization and integration of MULE-like transposons, another, possibly stress-activated, mechanism that can contribute to genetic and phenotypic diversity at this locus and across the *Phaeodactylum* genome, as previously hypothesized (Maumus *et al*., 2009). Our findings add relevant insights into the dynamics driving genome evolution in diatoms, suggesting that TEs can be rapidly mobilized and play a central role in enriching diatoms’ genetic diversity. Our findings open intriguing new questions on the mechanism of activation of these genetic elements, such as whether their mobilization is part of a regulated mechanism triggered by environmental stress, or whether their activity is determined by the de-repression of a silencing mechanism. In addition, it remains to be tested whether TE integration occurs randomly or if it is targeted towards some specific recombination *hot-spots* in the genome, characterized by genetic or epigenetic factors. More broadly, since TEs compose a large part of all sequenced diatom genomes (Maumus *et al*., 2009), our finding underscore the relevance that these interchromosomal non-homologous re-arrangements have in the dynamic genome of diatoms. These, not only are involved in re-shuffling endogenous genes, but potentially also in incorporating external DNA during the mobilization process, possibly facilitating horizontal gene transfers, which in diatoms are particularly frequent (Bowler et al., 2008), as well as providing new micro-homology regions that could promote recombination events (Bourque *et al*., 2018).

Besides their central role in aquatic ecology, diatoms are also emerging microbial hosts for biotechnology and synthetic biology applications, and many genetic tools including gene knock-out, knock-down and overexpression, as well as genome editing, are now part of routine functional genetics experimental practices to decipher diatom gene functions. In all these applications, a critical aspect is the genetic stability of the transformed cell lines. At the same time, all these applications involve a selection step, usually involving antibiotics or cytotoxic agents. Our findings show that these stressful environmental conditions caused by the accumulation of toxic compounds or by-products could mobilize TEs. Selection agents such as 5-FOA, zeocin, and bleomycin, or others derived from the accumulation of heterologous products or deriving from industrial cultivation settings, may affect loci that regard the introduced artificial genetic constructs, but also other regions of the genome, generating different genotypes in the resulting transformant lines which usually go unnoticed. As a direct consequence of the findings of this work, it is evident that the *PtUMPS* locus has important limitations as auxotrophic selectable marker for genetic engineering and genome editing approaches in diatoms, as it can be targeted by both TE-mediated and TE-independent mutations.

On the other hand, these results also provide inspiring new avenues for strain engineering strategies. For example, if controllable through environmental conditions and coupled with suitable selection or reporter systems, the mobilization of MULE TEs could be exploited to enrich genetic variation in the process of improving diatom strains, as inducible mutagenesis system.

## Conclusions

By subjecting the *PtUMPS* genomic locus of *P. tricornutum* cultures to a strong selective pressure with 5-FOA at concentrations routinely employed for genetic manipulation experiments, we observed the mobilization of a novel MULE transposon within the timeframe of a single experiment across four independent cell lines, which conferred the same 5-FOA resistant phenotype. In addition, we also observed a TE-independent deletion to occur in a fifth cell line. Our findings represent the first time in which the mobilization of a transposable element in diatoms is observed and tracked in *“real time”* during a specific experiment and suggest that this may be triggered by stressful environmental conditions. Moreover, our findings suggest that the TE that we identified might employ a novel mobilization mechanism which might involve protein degradation, through the presence of diatom-specific E3 ubiquitin ligases. Our findings have relevant implications in understanding the mechanisms underpinning the mobilization of transposable elements, in their role in the biology and evolution of diatoms, as well as in their biotechnological applications.

## Experimental procedures

### Microbial strains and growth conditions

Axenic cultures of *Phaeodactylum tricornutum* CCAP1055/1 were grown in liquid ESAW medium (Berges *et al*., 2001). Where appropriate, *P. tricornutum* was cultured in ESAW containing 300 μg/mL 5-FOA (Sigma Aldrich, Australia) and 50 μg/mL uracil (Sigma Aldrich, Australia), or ESAW supplemented with 50 μg/mL zeocin (Invivogen, San Diego, CA, USA). Diatoms were cultured in fully controlled shaking incubators (Kuhner, Switzerland) under 100 μE m^-2^ s^-1^ light in 21 °C shaking at 95 rpm.

### Flow cytometry

All cell counts were performed by flow cytometry, using a BD CytoFLEX S flow cytometer (BD Biosciences). The cells were analysed at medium speed until 20,000 events were counted. FITC fluorescence was excited with a 488 nm laser and emission was acquired using a 525/40 nm optical filter.

### PCR based analyses

PCR amplification was performed using Q5 high fidelity polymerase (New England Biolabs, Hitchin, UK) and PCR screening was performed using GoTaq Flexi DNA polymerase (Promega, Wisconsin, United States) according to the manufacturer’s instructions. For high throughput PCR screening, single colonies were grown in 200 μl ESAW supplemented with relevant compounds for 10 days to increase biomass. A list of primers used can be found in Table S1.

### High molecular weight genomic DNA (gDNA) extraction, sequencing and genome assembly

DNA was extracted and sequenced as described in (George *et al*., 2020). Individual sample sequencing runs were stopped after an estimated minimum of 11 Gbs were sequenced, with the exception of sample WRT2 and WT. For sample WRT2 coverage estimation during sequencing was inconsistent with the actual coverage after basecalling, resulting in approximately half the estimated coverage. The sequencing run for the wild type sample was performed last and only stopped once the flow cell was saturated resulting in more than 2 Gbs. After each sequencing run was stopped the flow cell was washed using wash kit EXP-WSH004 (Oxford Nanopore) according to manufacturer’s specifications and re-loaded with a new sequencing library.

Basecalling was performed using Oxford Nanopore’s basecaller Guppy v4.0.15 using the high accuracy model r9.4.1_450bps_hac. Basecalling resulted in 678,874 Oxford Nanopore long-reads ranging from 65,123 (WTR3) to 212,812 (wild type control). Coverage of the samples range from ~18X for sample WTR3 to ~68X for the WT control (Suppl Table 5). Basecalled reads were assembled using Flye v2.8.1 (Kolmogorov *et al*., 2019) and error corrected with three rounds of Racon v1.4.17 (Vaser *et al*., 2017). Mapping of long reads for error correction was performed using Minimap2 v2.17 (Li, 2018, p.2). Final assemblies ranged from 33,260,708 nt for sample WTR3 to 35,990,700 nt in the WT control (Suppl. Table 6). We obtained 678,874 Oxford Nanopore long-reads ranging from 65,123 (WTR3) to 212,812 (wild type control) resulting in a per-sample sequencing coverage ranging from ~18X for sample WTR3 to ~68X for the WT control (Table S5). After quality control and error correction final Flye v2.8.1 (Kolmogorov *et al*., 2019) assemblies ranged from 33,260,708 nt for sample WTR3 to 35,990,700 nt in the WT control (Table S6).

### Comparative genomics and TE annotation

All final assemblies were mapped to the published *P. tricornutum* genome (https://protists.ensembl.org/Phaeodactylum_tricornutum) using Minimap2 v2.17. Resulting sam files were sorted, converted to bam format, and indexed using samtools v1.10 (Li *et al*., 2009). The *P. tricornutum* UPMS region was screened for Insertions and deletions using the Integrated Genomics Viewer (IGV) v2.10 (Robinson *et al*., 2011) and the indexed bam files. Sequences of potential insertions at the UMPS locus were extracted using IGV and translated into their reverse complements online (https://www.bioinformatics.org/sms/rev_comp.html). Sequence alignments and visualization was done using AliView v1.28 (Larsson, 2014) and the integrated aligner Muscle v3.8.31 (Edgar, 2004). The insertions/potential TEs in the UMPS region of the genome assemblies were confirmed as true insertions using Sniflles v1.0.12 (Sedlazeck *et al*., 2018). Read mapping for Sniffles was performed with Minimap2 v2.17 and resulting sam files sorted, converted to bam and indexed using samtools v1.10.

Gene prediction and functional annotation of the coding sequences in the insertions were performed using InterproScan v5.54.87 (Jones *et al*., 2014).

For validation of the expression of predicted coding sequences of the insertion publicly available *P. tricornutum* RNASeq datasets were downloaded from the Sequence Read Archive (SRA) at the National Center for Biotechnology information (NCBI) and aligned to the complete transposon sequence of sample WTR1 using the splice-aware aligner STAR v2.7.10a (Dobin *et al*., 2013). To validate the expression of the transposase and other putative coding sequences of the transposon publicly available *P. tricornutum* RNASeq datasets were downloaded from the Sequence Read Archive (SRA) at the National Center for Biotechnology information (NCBI) and aligned to the complete transposon sequence of sample WTR1 using the splice-aware aligner STAR v2.7.10a (Dobin *et al*., 2013).

### Data statement and accession numbers

Basecalled sequencing files for each sample in fastq format are available at the SRA under bioproject PRJNA873859. Error corrected genome assemblies for all isolates, Sniffles variant calls in vcf format, InterProScan annotations and fasta files of the complete TE sequences were deposited in an Open Science Forum repository at https://osf.io/7wtds/. Name mapping between is as follows: WTR1 = WTR_1_1, WTR2 = WTR_2_1, WTR3 = WTR_2_2, WTR4 = WTR_1_3, and WTR5 = WTR_1_4.

## Abbreviations

UMPS: uridine monophosphate synthase
5-FOA: 5-fluoroorotic acid
TE: transposable element
LTR: long terminal repats
TSD: target site duplications
MULE: Mutator-like-Element

## Acknowledgments

We recognize that this work was conducted on the lands of the Gadigal People of the Eora Nation. This work was supported by the University of Technology Sydney (UTS) and the CSIRO Synthetic Biology Future Science Platform. MF was supported by a CSIRO Synthetic Biology Future Science Platform, co-funded by CSIRO and UTS. Basecalling and bioinformatics analysis were facilitated by Blue Mountains Analytics, Springwood (Australia).

## Authors’ contributions

JG designed the study, conducted the experiments, analysed data and wrote the manuscript. TK designed study, performed DNA sequencing, analysed data and contributed writing the manuscript; AC performed DNA extraction; RA analysed data and contributed writing the manuscript; MF designed the study, analysed data, and wrote the manuscript. All authors read and edited the manuscript.

## References

Armbrust, E.V. (2009) The life of diatoms in the world’s oceans. Nature, 459, 185–92.

Backhaus, K., Ludwig-Radtke, L., Xie, X. and Li, S.M. (2017) Manipulation of the Precursor Supply in Yeast Significantly Enhances the Accumulation of Prenylated β-Carbolines. ACS Synthetic Biology, 6, 1056–1064.

Berges, J.A., Franklin, D.J. and Harrison, P.J. (2001) EVOLUTION OF AN ARTIFICIAL SEAWATER MEDIUM: IMPROVEMENTS IN ENRICHED SEAWATER, ARTIFICIAL WATER OVER THE LAST TWO DECADES. Journal of Phycology, 37, 1138–1145.

Bourque, G., Burns, K.H., Gehring, M., et al. (2018) Ten things you should know about transposable elements. Genome Biology, 19, 1–12.

Bowler, C., Allen, A.E., Badger, J.H., et al. (2008) The Phaeodactylum genome reveals the evolutionary history of diatom genomes. Nature, 456, 239–44.

Brandsen, B.M., Mattheisen, J.M., Noel, T. and Fields, S. (2018) A Biosensor Strategy for E. coli Based on Ligand-Dependent Stabilization. ACS Synthetic Biology, 7, 1990–1999.

Bulankova, P., Sekulić, M., Jallet, D., et al. (2021) Mitotic recombination between homologous chromosomes drives genomic diversity in diatoms. Current Biology, 1–12.

Butler, T., Kapoore, R.V. and Vaidyanathan, S. (2020) Phaeodactylum tricornutum: A Diatom Cell Factory. Trends in Biotechnology, 1–17.

D’Adamo, S., Schiano di Visconte, G., Lowe, G., Szaub-Newton, J., Beacham, T., Landels, A., Allen, M.J., Spicer, A. and Matthijs, M. (2018) Engineering the unicellular alga *Phaeodactylum tricornutum* for high-value plant triterpenoid production. Plant Biotechnology Journal, 1–13.

Dobin, A., Davis, C.A., Schlesinger, F., Drenkow, J., Zaleski, C., Jha, S., Batut, P., Chaisson, M. and Gingeras, T.R. (2013) STAR: ultrafast universal RNA-seq aligner. Bioinformatics, 29, 15–21.

Dupeyron, M., Singh, K.S., Bass, C. and Hayward, A. (2019) Evolution of Mutator transposable elements across eukaryotic diversity. Mobile DNA, 10.

Edgar, R.C. (2004) MUSCLE: multiple sequence alignment with high accuracy and high throughput. Nucleic Acids Research, 32, 1792–1797.

Fabris, M., Abbriano, R.M., Pernice, M., et al. (2020) Emerging Technologies in Algal Biotechnology: Toward the Establishment of a Sustainable, Algae-Based Bioeconomy. Frontiers in Plant Science, 11, 1–22.

Fabris, M., George, J., Kuzhiumparambil, U., Lawson, C.A., Jaramillo Madrid, A.C., Abbriano, R.M., Vickers, C.E. and Ralph, P. (2020) Extrachromosomal genetic engineering of the marine diatom Phaeodactylum tricornutum enables the heterologous production of monoterpenoids. ACS synthetic biology.

Filloramo, G.V., Curtis, B.A., Blanche, E. and Archibald, J.M. (2021) Re-examination of two diatom reference genomes using long-read sequencing., 1–25.

George, J., Kahlke, T., Abbriano, R.M., Kuzhiumparambil, U., Peter, R. and Fabris, M. (2020) Metabolic engineering strategies in diatoms reveal unique phenotypes and genetic configurations with implications for algal genetics and synthetic biology. Frontiers in Bioengineering and Biotechnology, 8, 513.

Hanada, K., Vallejo, V., Nobuta, K., Slotkin, R.K., Lisch, D., Meyers, B.C., Shiu, S.-H. and Jiang, N. (2009) The Functional Role of Pack-MULEs in Rice Inferred from Purifying Selection and Expression Profile. The Plant Cell, 21, 25–38.

Hao, H., Wang, X., Jia, H., Yu, M., Zhang, X., Tang, H. and Zhang, L. (2016) Large fragment deletion using a CRISPR/Cas9 system in Saccharomyces cerevisiae. Analytical Biochemistry, 509, 118–123.

Hempel, F., Bozarth, A.S., Lindenkamp, N., Klingl, A., Zauner, S., Linne, U., Steinbüchel, A. and Maier, U.G. (2011) Microalgae as bioreactors for bioplastic production. Microbial cell factories, 10, 81.

Hempel, F., Lau, J., Klingl, A. and Maier, U.G. (2011) Algae as protein factories: Expression of a human antibody and the respective antigen in the diatom Phaeodactylum tricornutum. PLoS ONE, 6, e28424.

Huerta-Cepas, J., Szklarczyk, D., Heller, D., et al. (2019) eggNOG 5.0: a hierarchical, functionally and phylogenetically annotated orthology resource based on 5090 organisms and 2502 viruses. Nucleic Acids Research, 47, D309–D314.

Ishii, Y., Maruyama, S., Fujimura-Kamada, K., Kutsuna, N., Takahashi, S., Kawata, M. and Minagawa, J. (2018) Isolation of uracil auxotroph mutants of coral symbiont alga for symbiosis studies. Scientific Reports, 8, 1–9.

Jones, P., Binns, D., Chang, H.-Y., et al. (2014) InterProScan 5: genome-scale protein function classification. Bioinformatics, 30, 1236–1240.

Kabelitz, T., Brzezinka, K., Friedrich, T., Górka, M., Graf, A., Kappel, C. and Bäurle, I. (2016) A JUMONJI protein with E3 ligase and histone H3 binding activities affects transposon silencing in Arabidopsis. Plant Physiology, 171, 344–358.

Klein, J., Heal, J.R., Hamilton, W.D.O., Boussemghoune, T., Tange, T.Ø., Delegrange, F., Jaeschke, G., Hatsch, A. and Heim, J. (2014) Yeast synthetic biology platform generates novel chemical structures as scaffolds for drug discovery. ACS Synthetic Biology, 3, 314–323.

Kolmogorov, M., Yuan, J., Lin, Y. and Pevzner, P.A. (2019) Assembly of long, error-prone reads using repeat graphs. Nat Biotechnol, 37, 540–546.

Kondo, T., Johnson, C.H. and Hastings, J.W. (1991) Action spectrum for resetting the circadian phototaxis rhythm in the CW15 strain of Chlamydomonas: I. Cells in darkness. Plant Physiology, 95, 197–205.

Larsson, A. (2014) AliView: a fast and lightweight alignment viewer and editor for large datasets. Bioinformatics, 30, 3276–3278.

Li, H. (2018) Minimap2: pairwise alignment for nucleotide sequences. Bioinformatics, 34, 3094–3100.

Lv, Y., Edwards, H., Zhou, J. and Xu, P. (2019) Combining 26s rDNA and the Cre-loxP System for Iterative Gene Integration and Efficient Marker Curation in Yarrowia lipolytica. ACS Synthetic Biology, 8, 568–576.

Maclennan, M., García-Cañadas, M., Reichmann, J., et al. (2017) Mobilization of LINE-1 retrotransposons is restricted by Tex19.1 in mouse embryonic stem cells. eLife, 6, 1–32.

Maumus, F., Allen, A.E., Mhiri, C., Hu, H., Jabbari, K., Vardi, A., Grandbastien, M.A. and Bowler, C. (2009) Potential impact of stress activated retrotransposons on genome evolution in a marine diatom. BMC Genomics, 10, 1–19.

Minoda, A., Sakagami, R., Yagisawa, F., Kuroiwa, T. and Tanaka, K. (2004) Improvement of Culture Conditions and Evidence for Nuclear Transformation by Homologous Recombination in a Red Alga, Cyanidioschyzon merolae 10D. Plant and Cell Physiology, 45, 667–671.

Pampuch, M., Walker, E.J.L. and Karas, B.J. (2022) Towards synthetic diatoms: The Phaeodactylum tricornutum Pt-syn 1.0 project. Current Opinion in Green and Sustainable Chemistry, 35, 100611.

Robinson, J.T., Thorvaldsdóttir, H., Winckler, W., Guttman, M., Lander, E.S., Getz, G. and Mesirov, J.P. (2011) Integrative genomics viewer. Nat Biotechnol, 29, 24–26.

Sakaguchi, T., Nakajima, K. and Matsuda, Y. (2011) Identification of the UMP synthase gene by establishment of Uracil auxotrophic mutants and the phenotypic complementation system in the marine diatom Phaeodactylum tricornutum. Plant Physiology, 156, 78–89.

Scott, L.H., Mathews, J.C., Flematti, G.R., Filipovska, A. and Rackham, O. (2018) An Artificial Yeast Genetic Circuit Enables Deep Mutational Scanning of an Antimicrobial Resistance Protein. ACS Synthetic Biology, 7, 1907–1917.

Sedlazeck, F.J., Rescheneder, P., Smolka, M., Fang, H., Nattestad, M., Haeseler, A. von and Schatz, M.C. (2018) Accurate detection of complex structural variations using single-molecule sequencing. Nat Methods, 15, 461–468.

Serif, M., Dubois, G., Finoux, A.-L., Teste, M.-A., Jallet, D. and Daboussi, F. (2018) One-step generation of multiple gene knock-outs in the diatom Phaeodactylum tricornutum by DNA-free genome editing. Nature Communications, 9, 3924.

Slattery, S.S., Diamond, A., Wang, H., et al. (2018) An Expanded Plasmid-Based Genetic Toolbox Enables Cas9 Genome Editing and Stable Maintenance of Synthetic Pathways in Phaeodactylum tricornutum. ACS Synthetic Biology, 7, 328–338.

Slattery, S.S., Giguere, D.J., Stuckless, E.E., et al. (2022) Phosphate-regulated expression of the SARS-CoV-2 receptor-binding domain in the diatom Phaeodactylum tricornutum for pandemic diagnostics. Sci Rep, 12, 7010.

Standage-Beier, K., Zhang, Q. and Wang, X. (2015) Targeted Large-Scale Deletion of Bacterial Genomes Using CRISPR-Nickases. ACS Synthetic Biology, 4, 1217–1225.

Sun, L., Jing, Y., Liu, X., Li, Q., Xue, Z., Cheng, Z., Wang, D., He, H. and Qian, W. (2020) Heat stress-induced transposon activation correlates with 3D chromatin organization rearrangement in Arabidopsis. Nature Communications, 11.

Vancaester, E., Depuydt, T., Osuna-cruz, C.M. and Vandepoele, K. (2020) Systematic and functional analysis of horizontal gene transfer events in diatoms.

Vandepoele, K., Van Bel, M., Richard, G., Van Landeghem, S., Verhelst, B., Moreau, H., Van de Peer, Y., Grimsley, N. and Piganeau, G. (2013) pico-PLAZA, a genome database of microbial photosynthetic eukaryotes. Environmental microbiology, 15, 2147–53.

Vaser, R., Sović, I., Nagarajan, N. and Šikić, M. (2017) Fast and accurate de novo genome assembly from long uncorrected reads. Genome Res., 27, 737–746.

Wang, Y.K., Das, B., Huber, D.H., Wellington, M., Kabir, M.A., Sherman, F. and Rustchenko, E. (2004) Role of the 14-3-3 protein in carbon metabolism of the pathogenic yeast Candida albicans. Yeast, 21, 685–702.

Wellington, M., Kabir, M.A. and Rustchenko, E. (2006) 5-Fluoro-orotic acid induces chromosome alterations in genetically manipulated strains of Candida albicans. Mycologia, 98, 393–398.

Wellington, M. and Rustchenko, E. (2005) 5-Fluoro-orotic acid induces chromosome alterations in Candida albicans. Yeast, 22, 57–70.

Wicker, T., Sabot, F., Hua-van, A., et al. (1989) TEs_Classification_2007. Available at: https://search-proquest-com.dbgw.lis.curtin.edu.au/docview/223751027?OpenUrlRefId=info:xri/sid:primo&accountid=10382.

Windhagauer, M., Abbriano, R.M., Pittrich, D.A. and Doblin, M.A. (2022) Phosphate-inducible poly-hydroxy butyrate production dynamics in CO2 supplemented upscaled cultivation of engineered Phaeodactylum tricornutum. J Appl Phycol. Available at: https://link.springer.com/10.1007/s10811-022-02795-y [Accessed July 19, 2022].

