## Supporting Information for "Real time mobilization of a novel diatom *Mutator-Like Element* (MULE) transposon to inactivate the uridine monophosphate synthase (UMPS) locus in *Phaeodactylum tricornutum*"

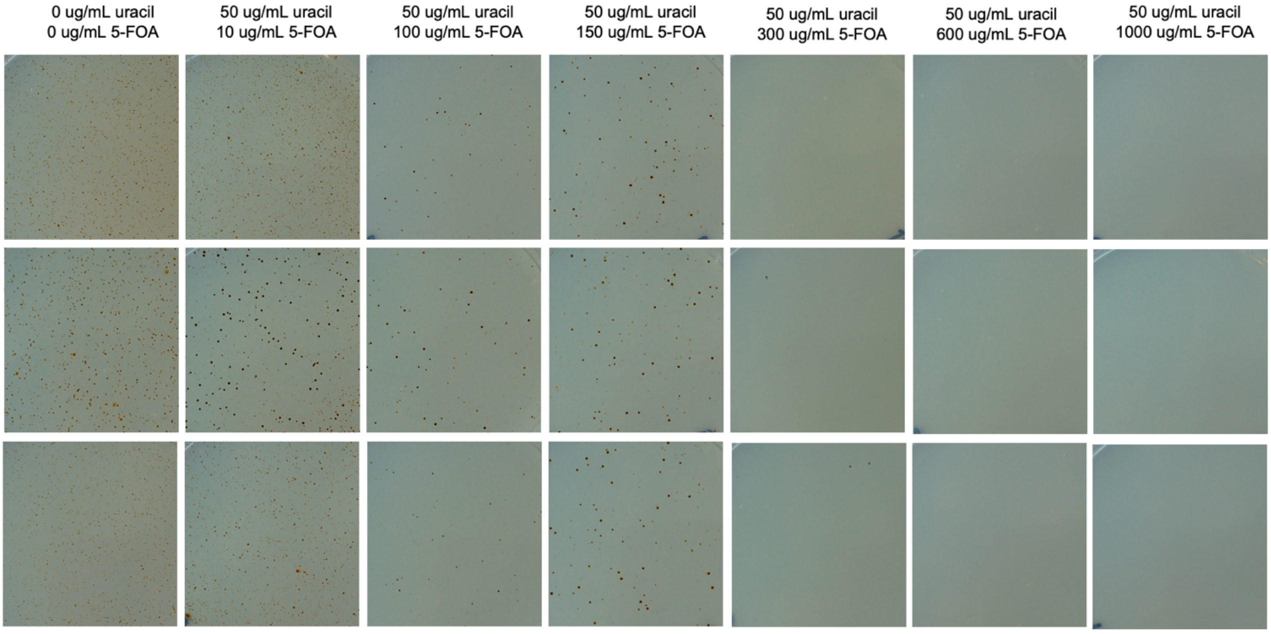

**Figure S1.** Spontaneous formation of *P. tricornutum* colonies on solid ESAW medium, and on solid ESAW medium with increasing concentrations of 5-FOA (10-1000 µg/ml) and supplemented uracil (50 µg/ml). Each panel (n=3) is a representative area on the same size for each plate.

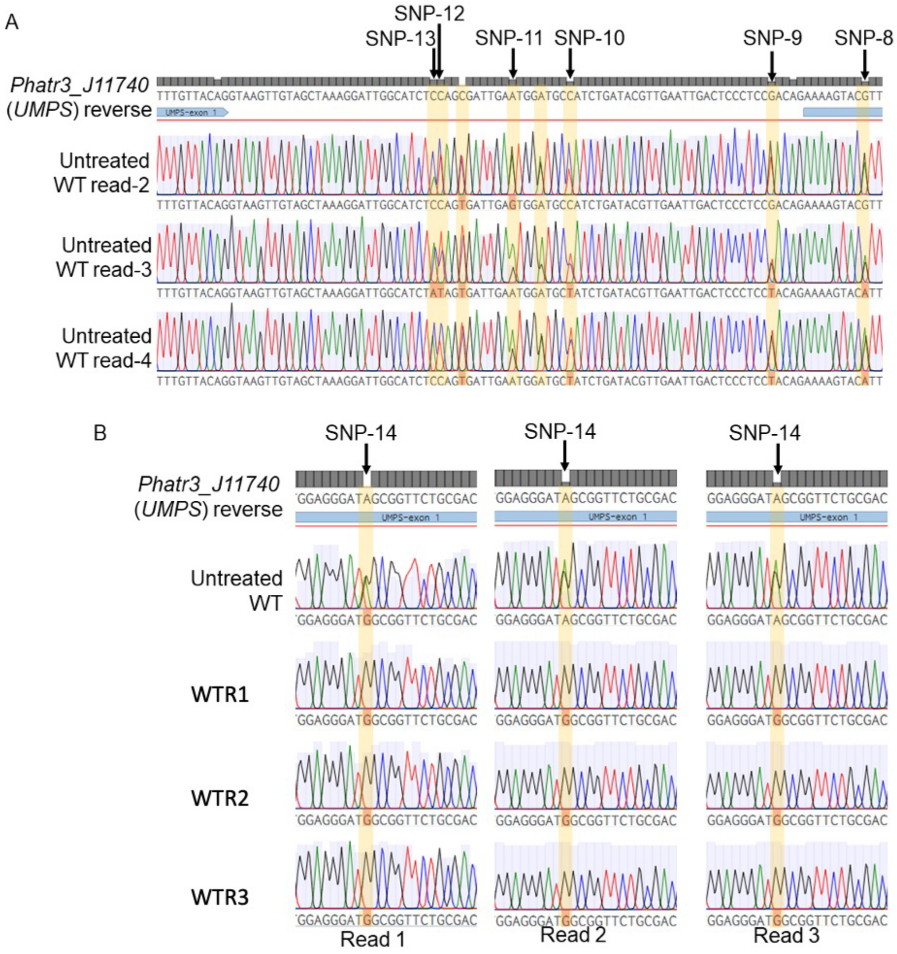

**Figure S2.** Analysis of *UMPS* genotype in untreated wild type *P. tricornutum* CCAP 1055/1 strain (A) The presence of doublet peaks at precise locations indicate single nucleotide polymorphisms characteristic of two non-identical gene copies (alleles) resulting in a heterozygous genotype in diploid organisms. These doublets occur in exogenic (blue) and intronic regions. The SNPs identified in this region are identical to those identified by Sakaguchi et al. (2011). (B) SNP-14 in wild type strain has been lost in all three WTR strains as evident by the loss of the doublet peak confirmed across three individually sequenced reads per cell line, indicating a homozygous genotype.

**Table S1.** Oligonucleotide primers used in this study.

| Primer ID | Sequence 5'-3' | Description |
|---|---|---|
| 40 | CAACAAAGTGCTCCTGCAAA | Forward primer to amplify *UMPS* from *Phatr3_45195* |
| 38 | ATGGCCACCCCCTCTTTTCGATCA | Forward primer to amplify *UMPS* from *UMPS* start codon |
| 75 | GAAGAAAATCGCTGTGACGC | Forward primer to amplify *UMPS* from within exon 1 of *UMPS;* to amplify *UMPS* for RE analysis |
| 74 | GTCCGTAGCTTTGCTGATACC | Reverse primer to amplify *UMPS* until *Phatr3_45193;* to amplify *UMPS* for RE analysis |
| 37 | TTACACTCCGTATTCGTTTCGAT | Reverse primer to amplify *UMPS* until intergenic region between *UMPS* and *Phatr3_45193* |
| 39 | CCTCTGCTTTCCGCATGTAT | Reverse primer to amplify *UMPS* until *UMPS* stop codon |
| MF941 | CCTCTGCTTTCCGCATGTAT | Sequencing *UMPS* amplicon |
| MF942 | AAGTGTGCGACTCACGAATG | Sequencing *UMPS* amplicon |
| MF943 | GAGCCGAATTTGAGAACACC | Sequencing *UMPS* amplicon |
| MF944 | ATCCAACAAAATCGGCACAT | Sequencing *UMPS* amplicon |
| MF945 | CAACACCAATTCGCTGATTC | Sequencing *UMPS* amplicon |
| MF946 | GCTCAGCAGACCGAGAGTTC | Sequencing *UMPS* amplicon |
| MF947 | AAGATTCCCGGATCAAACAA | Sequencing *UMPS* amplicon |

**Table S2.** Oxford Nanopore sequencing statistics

| Sample | #Reads (total) | Nucleotides (total) | N50 | Average read length | Coverage | # Inserts highly similar to the TE found in *PtUMPS* |
|---|---|---|---|---|---|---|
| WTR1 | 87,984 | 1,256,293,972 | 22,517 | 14,278 | ~35 | 118 |
| WTR2 | 99,286 | 1,156,842,678 | 18,607 | 11,651 | ~33 | 131 |
| WTR3 | 65,213 | 642,202,632 | 14,984 | 9,847 | ~18 | 20 |
| WTR4 | 115,726 | 1,100,879,520 | 16,153 | 9,512 | ~31 | 25 |
| WTR5 | 97,853 | 1,126,954,949 | 17,007 | 11,516 | ~32 | 109 |
| WT | 212,812 | 2,386,396,674 | 19,386 | 11,213 | ~68 | 5 |

**Table S3.** Genome assembly summary statistics

| Sample | # Nucleotides (total) | # Contigs | N50 |
|---|---|---|---|
| WTR1 | 35,918,073 | 408 | 180,383 |
| WTR2 | 35,138,244 | 2,020 | 172,022 |
| WTR3 | 33,260,708 | 1,679 | 232,543 |
| WTR4 | 34,902,037 | 235 | 139,996 |
| WTR5 | 34,884,087 | 733 | 164,265 |
| WT | 35,990,700 | 567 | 150,450 |

**Table S4.** Low GC, 9 bp target site duplication (TSD) sequences flanking transposon insertions at the *PtUMPS* locus. Orientation was arbitrarily assigned during genome assembly.

| Sample | Target site duplication | Orientation | Length (bp) | GC% |
|---|---|---|---|---|
| WTR1 | AAACGAGTT | + | 9 | 33.3 |
| WTR2 | TTTCTTTTT | - | 9 | 11.1 |
| WTR3 | TTTCTTTTT | - | 9 | 11.1 |
| WTR5 | TTTCTTGAA | - | 9 | 22.2 |

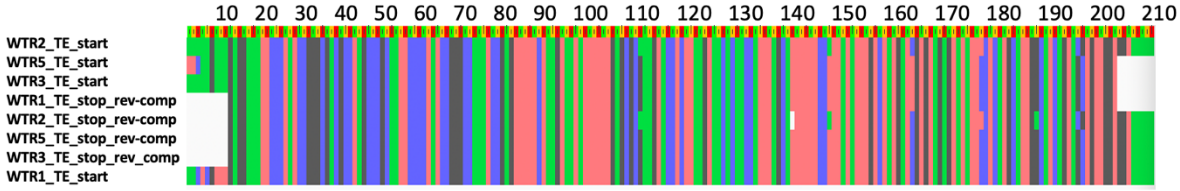

**Figure S3.** Alignments of the first 210 nucleotides of the TE sequence of samples WTR1, 2, 3, and 5 and the reverse-complement of the last 210 nucleotides showing the high similarity of the inverse terminal repeats of the TE sequences.

**Table S5.** BLAST analyses of the putative ORFs containing MULE transposase and E3 ligase component domains in TE sequences. Full putative ORFs found in WTR1 were used as queries, positions indicated are relative to ORFs in WTR1.

| **ORF containing MULE transposase domain, Zinc finger, SWIM-type domain** | | | |
|---|---|---|---|
| **Sample** | **start position** | **stop position** | **% identity and translated protein length** |
| **WTR1** | 320 | 2224 | full, used as query |
| **WTR2** | 1972 | 1847 | 69%, 2-43aa |
| **WTR2** | 1842 | 1792 | ~60%, ~44-60aa |
| **WTR2** | 1790 | 1467 | 89%, ~526-635aa |
| **WTR3** | 3397 | 1493 | 100%, full sequence |
| **WTR5** | 2869 | 3378 | 100%, WTR1 1-170aa |
| **WTR5** | 2612 | 2872 | 100%, WTR1 179-256aa |
| **WTR5** | 2337 | 2612 | 100%, WTR1 257-348aa |
| **WTR5** | 2143 | 2340 | 77%, WTR1 348-413aa |
| **WTR5** | 1481 | 2227 | 74%, WTR1 386-635aa |
| **Putative ubiquitin protein ligase E3, zinc finger UBR type** | | | |
| **Sample** | **start** | **stop** | **% identity and translated protein length** |
| **WTR1** | 2531 | 2959 | full, used as query |
| **WTR2** | 1189 | 764 | 68%, 1-143aa |
| **WTR3** | 1186 | 758 | 100%, 1-143aa |
| **WTR5** | 1177 | 800 | 53%, 1-135aa |
| **WTR5** | 884 | 759 | 100%, 102-143aa |
